# VIL1, a Polycomb-associated protein, modulates high ambient temperature response via H3K27me3 and H2A.Z in *Arabidopsis thaliana*

**DOI:** 10.1101/2020.04.29.069484

**Authors:** Yogendra Bordiya, Junghyun Kim, Yanpeng Xi, Dong-Hwan Kim, Youngjae Pyo, Bo Zhao, Wei Zong, William A. Ricci, Xiaoyu Zhang, Sibum Sung

## Abstract

Adapting to the everchanging environment is key to a successful life for an organism. Eukaryotes reprogram their transcriptome in order to adapt to an unfavorable environment. To achieve this reprogramming, plants and animals employ multiple responses including epigenetic regulation. In the search for mutations compromised in high ambient temperature response, we found that VIL1, a PHD finger protein displays aberrant development at high temperature. RNA-seq analysis shows that *vil1* fails to downregulate heat suppressed genes. H2A.Z ChIP-seq showed that unlike wild type, *vil1* fails to evict H2A.Z from heat responsive genes. We also found that *vil1* suppresses constitutive thermo-morphogenic phenotype of *arp6*. Supporting this phenotype, RNA-seq analysis revealed that constitutive heat responsive transcriptome of *arp6* reverted back to the wild-type levels in *arp6vil1*. This observation suggests an antagonistic relationship between VIL1 and ARP6. We found that this antagonism can be explained in part by interaction between H3K27me3 and H2A.Z.

## Introduction

Normal development is the first step towards completing a successful life cycle. The ability of organisms to grow, reproduce and achieve their maximum physical potential can be severely affected by the environmental perturbations. Changes in the climate are leading to widespread variation in important environmental factors such as temperature. The average global temperature on Earth has increased by about 0.6°C in the last century ^1^. Most of this warming has occurred in the past three decades at the rate of 0.15-0.20 °C per decade. Increase in global temperature caused by climate change will have a devastating effect on biodiversity, crop yield and ultimately on human health ^2-6^. Adapting to the everchanging environment is key to the successful life of an organism and ultimately of a species. To cope with unfavorable conditions, eukaryotes employ multiple responses including epigenetic regulation. These epigenetic regulations help in reprogramming of the transcriptome to cope with varying conditions. This process requires conversion of the environmental stimuli to a molecular signal. Much progress has been made to identify receptors to these sorts of environmental stimuli. Studies have also implicated the role of histone variant H2A.Z in mediating the thermo-sensory response in plants. An evolutionarily conserved SWR1 complex deposits histone variant H2A.Z and regulates multiple developmental processes in eukaryotes ^7^. Multiple reports have shown that H2A.Z is evicted from the histone octamer to be replaced with canonical H2A in response to high ambient temperature, however the molecular mechanism behind this eviction remains unknown.

To identify additional regulators of ambient temperature responses, we took reverse genetic approach to uncover novel epigenetic components involved in response to high ambient temperature. Namely, we screened a number of mutants previously shown to be involved in the chromatin modifications. Surprisingly, we identified that mutations in VIL1 (VIN3-LIKE 1)/VERNALIZATION 5(VRN5) impair high ambient temperature responses. VIL1, a PRC2 (Polycomb Repression Complex 2) associated protein, is known to mediate a cold-temperature response, vernalization ^8,9^. Previous studies also implicated the role of a component of the SWR1 complex in mediating temperature response ^10-16^. This prompted us to investigate the mechanisms by which temperature-triggered responses are mediated through both PRC2 and SWR1. We found that H2A.Z’s thermo-sensory response is influenced by the function of VIL1. Here, we show that VIL1 is required for response to warm ambient temperature because of two main factors: 1) by downregulating the heat suppressed genes through its PRC2 associated function, and; 2) by evicting H2A.Z in from heat responsive genes.

## Results

### *vil1* is hyposensitive to high ambient temperature

We took reverse genetic approach to uncover novel player in high ambient temperature in plants. For this purpose, we screened an array of mutants of genes ranging from histone readers, writers, erasers, chromatin remodelers to histone variants and found that mutation in VIL1, a PRC2 associated protein, displayed aberrant development at high ambient temperature of 27°C compared to WT (Fig. 1, Supplementary Fig S1). VIL1 is a PHD finger domain containing protein which has been shown to participate in both the photoperiod and vernalization flowering pathways in Arabidopsis by regulating expression of the related floral repressors *FLOWERING LOCUS C* and *FLOWERING LOCUS M* ^8^. PRC2 biochemically copurifies with members of the VERNALIZATION INSENSITIVE3 (VIN3) family of proteins, including VIN3, VIL1 and VIL2 ^17,18^. All VIN3 family proteins including VIL1 are known to act together with PRC2 to increase repressive histone marks at floral repressor loci in Arabidopsis ^18^. In order to adapt to the high temperature environment, plants undergo thermo-morphogenesis. Thermo-morphogenesis is morphological and architectural changes induced by high ambient temperatures ^19-21^. Thermo-morphogenesis at elevated ambient temperature involves changes in plant architecture such as elongation of hypocotyl in seedlings and extended petiole length and hyponasty in the rosette to keep plant body well above hot ground and facilitate aeration ^19^. We found that unlike WT, *vil1* does not show extended petiole length (Fig. 1A). Moreover, when switched from 22°C to 27°C growth condition, relative hypocotyl lengthening was reduced in *vil1* (Supp. Fig. 1B and C). It is very well established that thermo-morphogenic phenotype of long hypocotyl and petiole length is achieved with the help of auxin ^22-25^. Consistent with this, the inducibility of auxin responsive genes like *YUC8* and *IAA29* has been compromised in *vil1* (Supp Fig. 1D). This poor response in terms of phenotype suggests that *vil1* is hyposensitive to warm temperature. We also checked the flowering time data and found that *vil1* displayed late flowering phenotype in short day condition compared to WT both at 22°C and 27°C (Fig. 2D).

**Figure 1.**
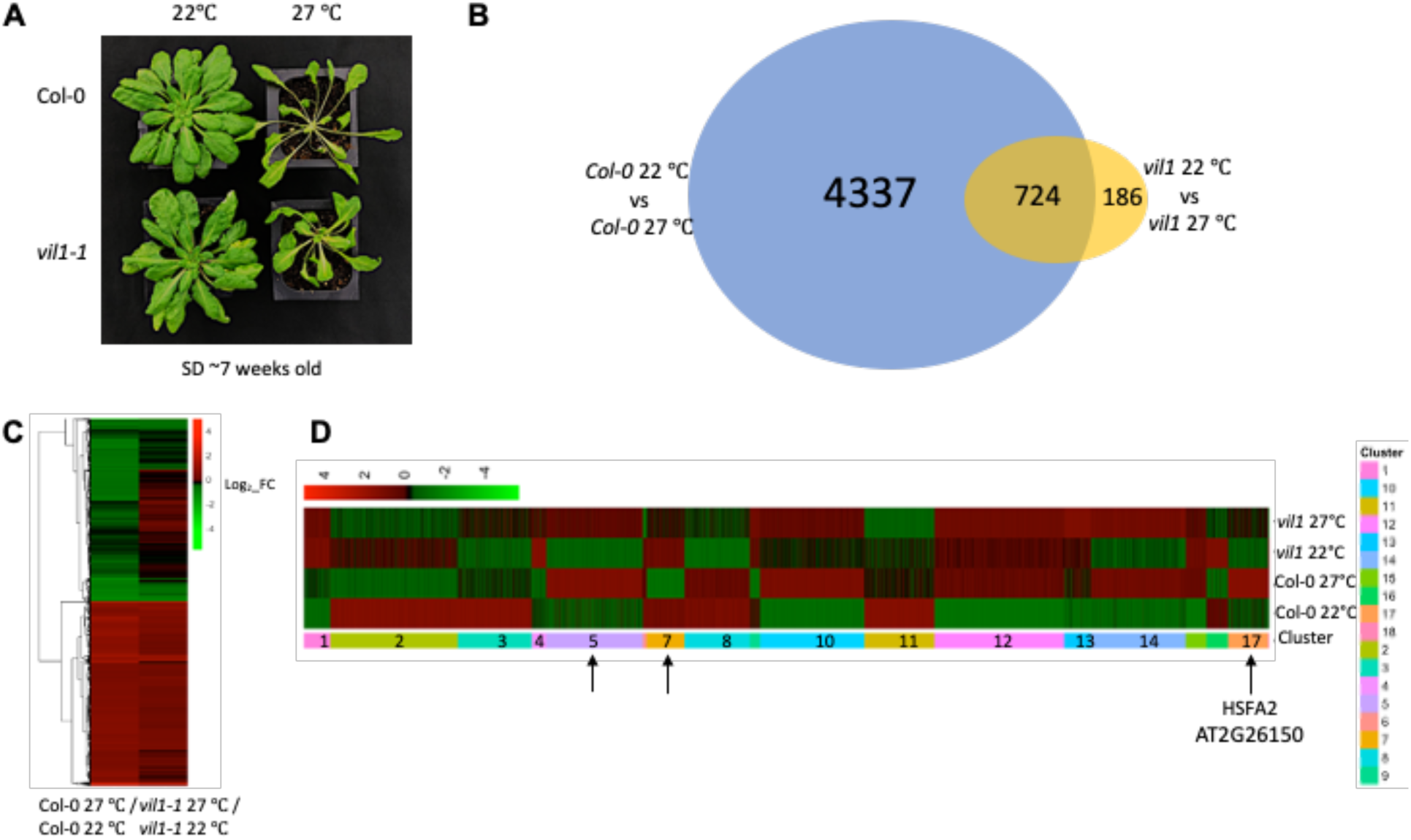
*vil1* responds poorly to high ambient temperature and fails to undergo thermo-morphogenesis. **A**. Col-0 (WT) and *vil1-1* plants grown in short day condition continuously at either 22°C or 27°C. As seen in the image, Col-0 undergo thermo-morphogenesis but *vil1-1* does not. **B**. Venn diagram overlap analysis of up and downregulated genes in response to high temperature between Col-0 and *vil1-1*. Number of genes respond to high temperature in *vil1-1* is much lower than in WT. **C**. Heatmap of up and downregulated genes show that *vil1-1* fails to downregulate majority heat suppressed genes and fails to upregulate a number of genes at high temperature compared to WT. **D**. Differentially expressed genes were used for hierarchical clustering analysis in order to find which genes are affected in *vil1-1*. Clusters 5, 7 and 17 indicated by an arrow contain genes which respond poorly to temperature change in *vil1-1*. Cluster 17 contains heat shock transcription factor 2. GO term analysis of these clusters is in supplementary figure 1.

**Figure 2.**
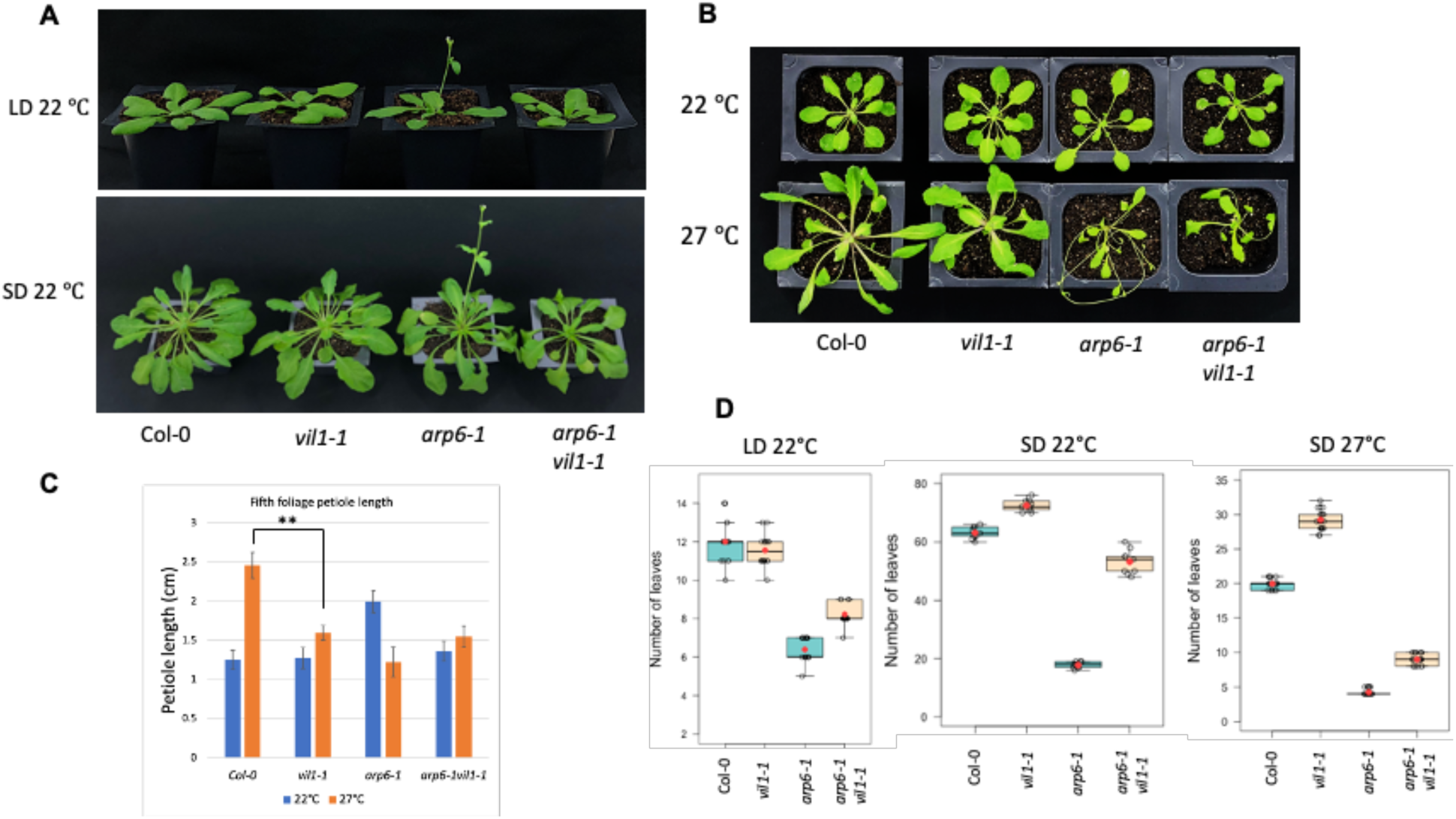
*vil1-1* suppresses constitutive thermo-morphogenic phenotype of *arp6-1*. **A**. Both in long day (LD) and in short day (SD) conditions, early flowering and long petiole length phenotype of *arp6* is suppressed by *vil1* in *arp6vil1*. **B and C**. phenotype at 22°C and 27°C and the petiole length quantification data. Petiole length of fifth leaf is shown. **D**. Flowering time data shown in terms of number of leaves at the time of flowering. Red dot in the box plot indicate mean number of leaves.

### Transcriptional changes upon high ambient temperature are compromised in *vil1* mutants

To address the molecular basis of hyposensitivity of *vil1* towards high temperature, we performed RNA-seq. First, we performed an overlap analysis between genes up and downregulated in WT and *vil1* at 22°C vs 27°C and found that there are 5,061 genes up- and down- regulated in response to high temperature in WT, whereas only 910 genes are up and down regulated in *vil1* (Fig. 1B). Heatmap analysis shows that *vil1* fails to downregulate majority of heat suppressed genes and also fails to upregulate a number of heat induced genes (Fig. 1C). In order to find specific genes responsible for hyposensitivity of *vil1* to high temperature, we performed hierarchical clustering analysis. Out of 17 clusters, clusters 5 and 17 contain genes which are not induced at high temperature as much as WT in *vil1* (Fig. 1D). GO (Gene Ontology) term analysis of these clusters showed that clusters 5 and 17 contain genes responsive to heat and stress (Supp Fig. 1A). Cluster 7, on the other hand contains genes which are suppressed by high temperature in WT but not in *vil1*. GO term analysis of cluster 7 shows enrichment of genes related to photosynthesis and response abiotic stimulus. It has been reported that photosynthetic activity is reduced at high ambient temperature in order to cope with the stress ^26^. These results also suggest that *vil1* fails to reduce photosynthetic activity at higher temperature unlike WT. RPKM normalized gene count from all RNA-seq samples is provided in supplementary table 3. Taken together, all these results suggest that *vil1* is hyposensitive in high ambient temperature in terms of morphological responses as well as of transcriptomic responses.

### *vil1* suppresses constitutive thermo-morphogenic phenotype of *arp6*

Switching from canonical histone to a non-canonical variant from histone octamer is a well-studied mode of chromatin remodeling ^27^. In Arabidopsis, a component of the SWR1 complex, ARP6 (Actin Related Protein 6), is involved in the deposition of histone variant H2A.Z ^28^. Loss of ARP6 in Arabidopsis leads to a major developmental and pleiotropic phenotype ^29^. H2A.Z has been implicated to play an important role in not only developmental regulation but also in the biotic and abiotic stress response ^10,30-32^. Recently, multiple reports implicated the role of H2A.Z in mediating thermo-sensory response in plants ^11-13^. Interestingly, previous studies have shown that the wild-type plants grown at higher ambient temperature (27°C) and *arp6* plants grown at normal ambient temperature (22°C) show a similar phenotype and transcriptome profile ^11^. This indicates that H2A.Z is a major regulator of the heat response in Arabidopsis. Remarkably, the PIF4 mediated thermo-morphogenesis requires removal of histone variant H2A.Z from the histone octamer at promoters of growth related genes ^14^. This also explains why *arp6* plants grown at 22°C phenocopy wild-type plants grown at 27°C.

It is known that under high ambient temperature and stress, plants accelerate the development and transition to flowering to be able to reproduce in time ^33^. *arp6* shows constitutive thermo-morphogenic phenotype and accelerated flowering (Fig. 2A) resulting from the constitutive expression of heat responsive genes ^11^. Consistently, *arp6* also shows longer petiole length at 22°C compared to WT (Fig. 2B and 2C). Strikingly, we found that *vil1* suppresses constitutive thermo-morphogenic phenotype of *arp6* suggesting that VIL1 and ARP6 have antagonistic functions (Fig. 2A and 2B). Unlike *arp6* single mutant, *arp6/vil1* double mutant no longer show long petiole length and early flowering phenotype associated with high ambient temperature (Fig. 2C and 2D). We confirmed this double mutant phenotype using another allele of *vil1* (*vil1-2*) (Supp. Fig. 5A).

### RNA-seq analysis supports suppression of thermo-morphogenic phenotype of *arp6-1/vil1- 1* double mutant and reveals antagonistic relationship between VIL1 and ARP6

In order to find out if the transcriptome supports antagonistic relationship between VIL1 and ARP6, we performed RNA-seq in Col-0 (WT), *vil1, arp6*, and *arp6vil1*. To confirm the previously published result that *arp6* at normal ambient temperature of 22°C displays a transcriptome similar to that of a plant grown at higher ambient temperature of 27°C ^11^, we performed a correlation analysis. For this purpose, we compared the transcriptome of WT plants grown at 27°C and *arp6* plants grown at 22°C. The result shows a clear positive correlation (Fig. 3A) confirming conclusion from previous studies. Note that all the transcribed genes in the genome were used for this correlation analysis. Since *vil1* is hyposensitive to high ambient temperature, when we compared the transcriptome of *vil1* plants grown at 27°C and *arp6* plants grown at 22°C, as expected, the positive correlation is lost to a great extent (Fig. 3B). Next, we took the similar approach considering only the genes which are up- and down-regulated in response to heat in WT and performed a boxplot analysis. This analysis elegantly captures the hyposensitivity of *vil1* and hypersensitivity of *arp6* to high temperature (Fig. 3C). Remarkably, exaggerated constitutive heat responsive transcriptome of *arp6* reverted back to the wild-type levels in *arp6vil1* (Fig. 3C). This result is consistent with the reversed morphological phenotype of *arp6vil1* compared to *arp6* which includes not only the change in flowering time but also unlike *arp6* single mutant, *arp6/vil1* double mutant no longer show long petiole length phenotype associated with high ambient temperature (Fig. 2A, 2B, 2C and 2D). We further confirmed this phenotype using another allele of *vil1* and via genetic complementation (Supp Fig. 5B). Next, to check if the reversal in the transcriptome in *arp6vil1* is limited to the heat responsive genes or to the overall transcriptome, we performed the correlation analysis of transcripts between WT and *arp6vil1* and found highly significant correlation (R^2^ = 0.85). This result suggest that majority of the transcripts differentially expressed in *arp6* compared to wild-type revert back to the wild-type level in *arp6vil1* (Fig. 3E). The top panel of supplementary figure 2 shows IGV screenshots of RNA-seq data showing reversion of transcriptome in *arp6vil1*. We confirmed the RNA-seq data with qRT-PCR on randomly selected genes on independent samples (Supp. Fig. 2). These results led us to the hypothesis that VIL1 and histone variant H2A.Z have antagonistic function in the regulation of gene expression in Arabidopsis.

**Figure 3.**
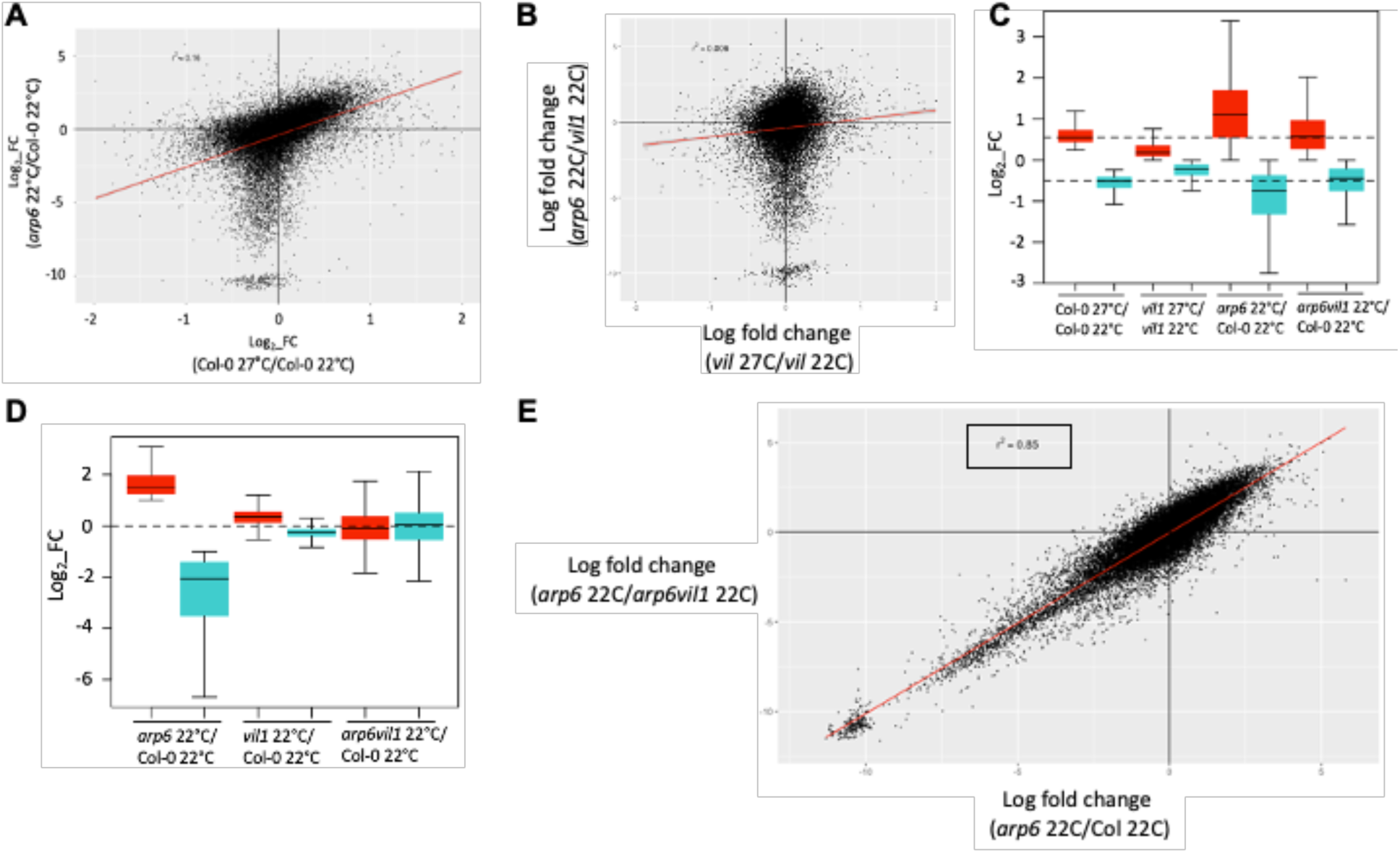
RNA-seq analysis supports suppression of thermo-morphogenic phenotype of *arp6-1/vil1-1* double mutant and reveals antagonistic relationship between VIL1 and ARP6. **A**. Correlation analysis using all actively transcribed genes in Arabidopsis. Comparison of *arp6* induced genome-wide transcriptional changes (*arp6* 22°C/Col-022°C) with responses to increasing temperature from 22°C to 27°C in wild-type shows that transcripts upregulated in arp6 mutant at 22°C positively correlated with transcripts induced by temperature. **B**. The correlation is lost in the same comparison where WT is replaced with vil1. **C**. Box plot was created using differentially expressed genes at 27°C compared to 22°C in WT. Red color indicates upregulated genes and cyan color shows downregulated genes. Notice that when same genes were taken from arp6 22C vs Col-0 22°C comparison, they shown constitutive differential expression on heat responsive genes. **D**. Genes differentially expressed in arp6 compared to WT (*arp6* 22°C/Col- 022°C) were used to create boxplot. These genes return to zero-fold change level in arp6vil1 22°C/Col-0 22°C comparison. **E**. All actively transcribed genes in Arabidopsis genome were used to create a scatter plot between the comparisons shown. Red line in all scatter plots represents linear regression line with r^2^ value of 0.85. FC (Fold Change).

### Clustering analysis reveal genes reversed in *arp6/vil1*

To find out how many and what kind of genes are reversed in *arp6vil1*, we performed hierarchical clustering of RNA-seq in Col-0 (wild-type), *vil1, arp6*, and *arp6vil1* followed by GO term analysis on these clusters. Out of ten clusters, clusters 1 and 2 with 3,797 and 3,840 genes respectively contain genes which are up and downregulated in *arp6* compared to the WT, however, their expression was back to the WT level in *arp6vil1* double mutant (Fig. 4A). The heatmap of these clusters is shown in Fig. 4B. In order to find out biological relevance behind this antagonism between VIL1 and ARP6 we performed GO term analysis on these clusters. No clear term to explain the relationship was found.

**Figure 4.**
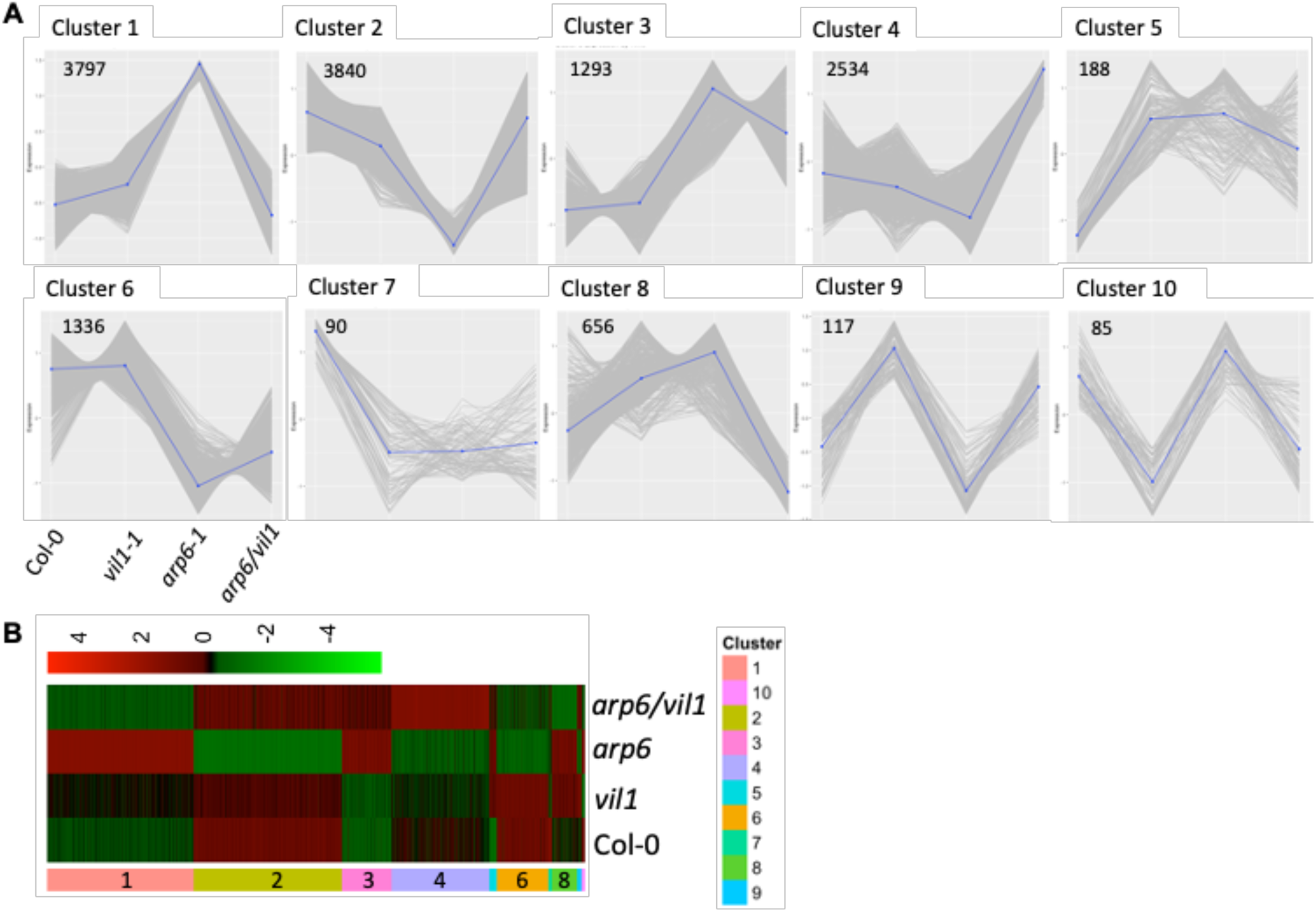
Clustering analysis reveal genes reversed in *arp6/vil1*. **A**. Differentially expressed genes were used for hierarchical clustering analysis in order to find which genes are transcriptionally reversed in *arp6vil1*. Out of 10 clusters, clusters 1 and 2 show transcriptionally reversed genes. **B**. Heatmap of 10 clusters.

**Figure 5.**
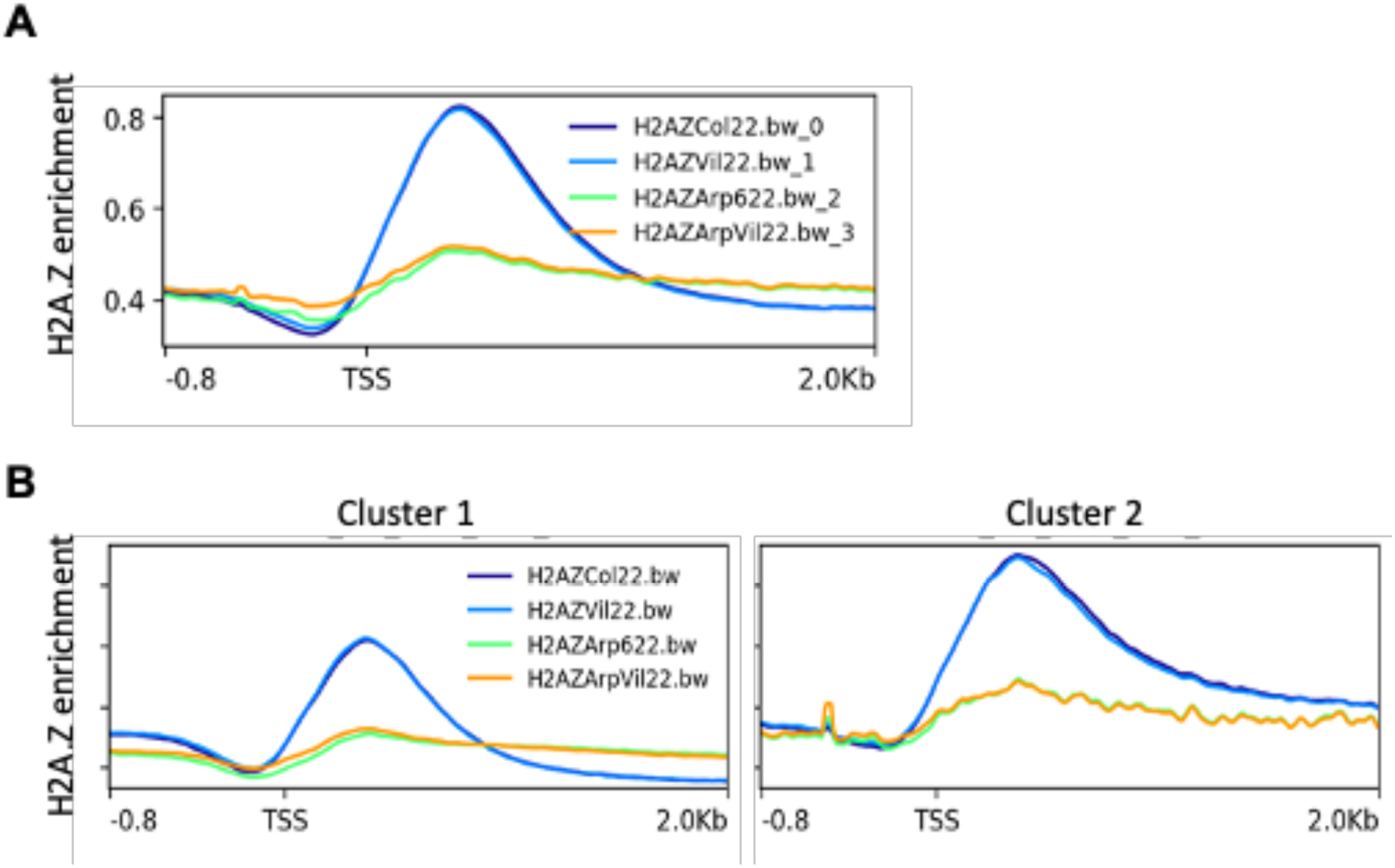
Overall H2A.Z level remain unchanged in *vil1-1* compared to WT. BED file was created using MACS2 generated H2A.Z ChIP peaks information. This BED file was fed into deepTools to generate compute matrix followed by profile plotting to find the enrichment of H2A.Z from 800 bp upstream of TSS to 2.0 kbp downstream of genes. **A**. Profile of all the genome peaks comparing H2A.Z enrichment among WT, *vil1, arp6* and *arp6vil1*. **B**. H2A.Z enrichment in genes from clusters 1 and 2 which are transcriptionally reversed in arp6vil1.

### Overall H2A.Z level remains unchanged in *vil1-1* compared to WT

We have two hypotheses to explain the antagonistic relationship between ARP6 and VIL1; 1) reduced deposition of H2A.Z in *arp6* and enhanced deposition of H2A.Z in *vil1* or: 2) Interaction between histone modifications, such as H3K27me3, and H2A.Z have antagonistic roles in regulation of gene expression. To check the first possibility, we performed the H2A.Z ChIP-seq at 22°C and in WT, *vil1, arp6* and *arp6vil1*. MACS2 ChIP-peak analysis revealed about 16,000 peaks in both WT and *vil1* (Supp Table 2) suggesting that there was no peak loss or gain in *vil1*. We then checked the relative enrichment of the H2A.Z using deepTools among four genotypes. As expected, the enrichment of H2A.Z was poor in *arp6* compared to the WT, however, there was no significant change between WT and *vil1* and between *arp6* and *arp6vil1* (Fig. 5A). To find out the basis of antagonism between ARP6 and VIL1, we did the H2A.Z enrichment analysis in clusters 1 and 2 which contain genes reversed in *arp6vil1*. We did not find significant difference between WT and *vil1* and between *arp6* and *arp6vil1* in both clusters (Fig. 5B). These results suggest that overall H2A.Z level remains unchanged in *vil1* compared to WT and that the antagonisms between VIL1 and ARP6 cannot be explained by H2A.Z enrichment alone.

### *vil1-1* fails to evict H2A.Z from high ambient temperature responsive genes

Given the role of H2A.Z in thermo-sensory response in Arabidopsis and genetic interaction between VIL1 and ARP6, we decided to perform H2A.Z ChIP-seq at 22°C and 27°C WT and *vil1*. As indicated in the previous section, at the global level, there was no significant change in the enrichment of H2A.Z between WT and *vil1* at 22°C. To look at the H2A.Z level in heat responsive genes specifically, we performed the differential binding analysis using Diffbind tool to compare the H2A.Z fold change between 22°C and 27°C. As reported in a number of studies, H2A.Z level reduced at 27°C compared to 22°C in WT (Fig. 6A), however, the level remained high in case of *vil1* at 27°C (Fig. 6B). To confirm that this change in level of H2A.Z at 27°C is correlated with change in gene expression, we looked at the RNA expression level change (Fig. 6C). We found that indeed the level of H2A.Z in heat responsive genes was reduced in WT and that this reduction was not seen in *vil1* leading to poor induction of heat responsive genes (Fig. 6C). When we looked at more heat responsive genes, similar pattern was found (Supp. Fig. 3). This result suggests that VIL1 is required for the eviction of H2A.Z from heat responsive genes. It also explains in part the hyposensitive nature of *vil1* mutants at high ambient temperature.

**Figure 6.**
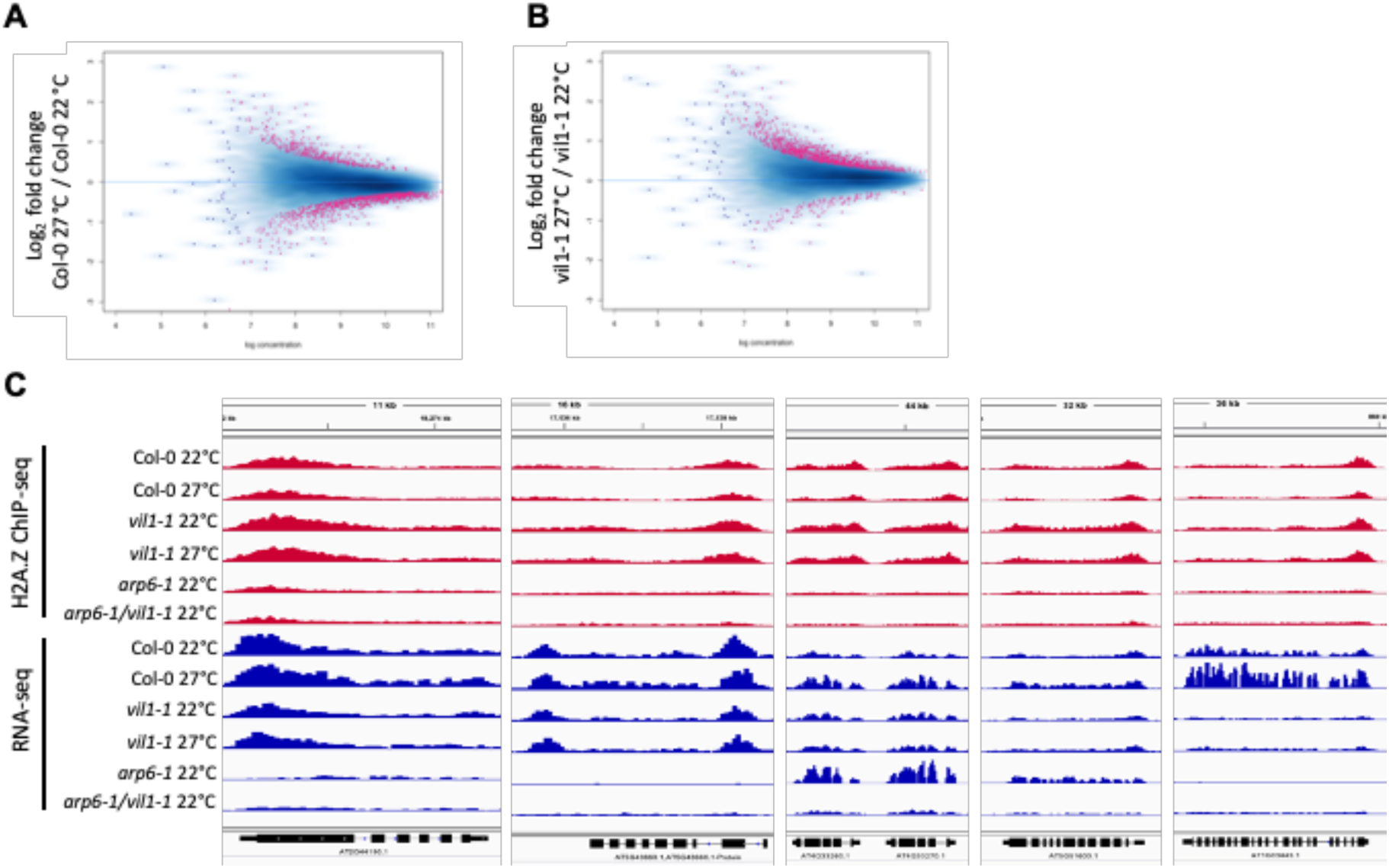
*vil1-1* fails to evict H2A.Z from high ambient temperature responsive genes. **A**. MA plot with log_2_ fold change in the H2A.Z enrichment. Differential enrichment in Col-0 27°C/Col-0 22°C comparison and **B**. in *vil1* 27°C/*vil1* 22°C comparison. **C**. IGV snapshots with H2A.Z ChIP-seq and RNA-seq data of randomly selected genes which fail to evict H2A.Z in *vil1* in response to high temperature.

### Antagonism between VIL1 and ARP6 in part can be explained by H3K27me3

We performed ChIP-qPCR on various histone modifications to check the possibility of interaction between H2A.Z and histone modifications. Studies suggest that the interaction between H2A.Z and PRC2 might be much more complex and involve additional histone modifications. Here we tried to examine relationship of H3K27me3, H3K4me2 and H3K4me3 and other modifications with H2A.Z. For this purpose, we randomly selected transcriptionally reversed genes in *arp6vil1* and performed ChIP-qPCR in WT, *vil1, arp6* and *arp6vil1* backgrounds. IGV snapshot of these genes is shown in Fig. 7A. We performed ChIP-qPCR to check H3K27me3, H3K9me2, H3K4me2, H3K4me3, H3 acetylation and RNA Pol II (S5). Interestingly, we did not find significant change in any of these histone modifications and RNA Pol II enrichment except for H3K27me3 (Fig. 7B and Supp Fig. 4). It’s clear that H3K27me3 level in low in *vil1* compared to WT which is expected given VIL1 is a PRC2 associated protein. Interestingly, the level of H3K27me3 was further reduced in *arp6vil1* double mutant. To gain more insight into the role of H3K27me3 in antagonism between VIL1 and ARP6, we performed overlap analysis between transcriptionally reversed genes in *arp6vil1* (clusters 1 and 2) and H3K27me3 enriched genes in Arabidopsis genome. 27.2% (1,045) genes from cluster 2 were enriched with H3K27me3 as opposed to mere 5.4% from cluster 1 (Fig. 7C). When we looked at the proportion of 1045 genes up and downregulated in *vil1, arp6*, and *arp6vil1* compared to WT, while all of these genes were downregulated in arp6, about 75% reversed back to upregulation in *arp6vil1* (Fig. 7D). To check if VIL1 binds at these 1045 reversed loci, we randomly selected 7 genes for VIL1 ChIP-qPCR. Except for AT4G29100 which was used as negative control, all other loci show VIL1 enrichment including FLC (Supp. Fig. 7).

**Figure 7.**
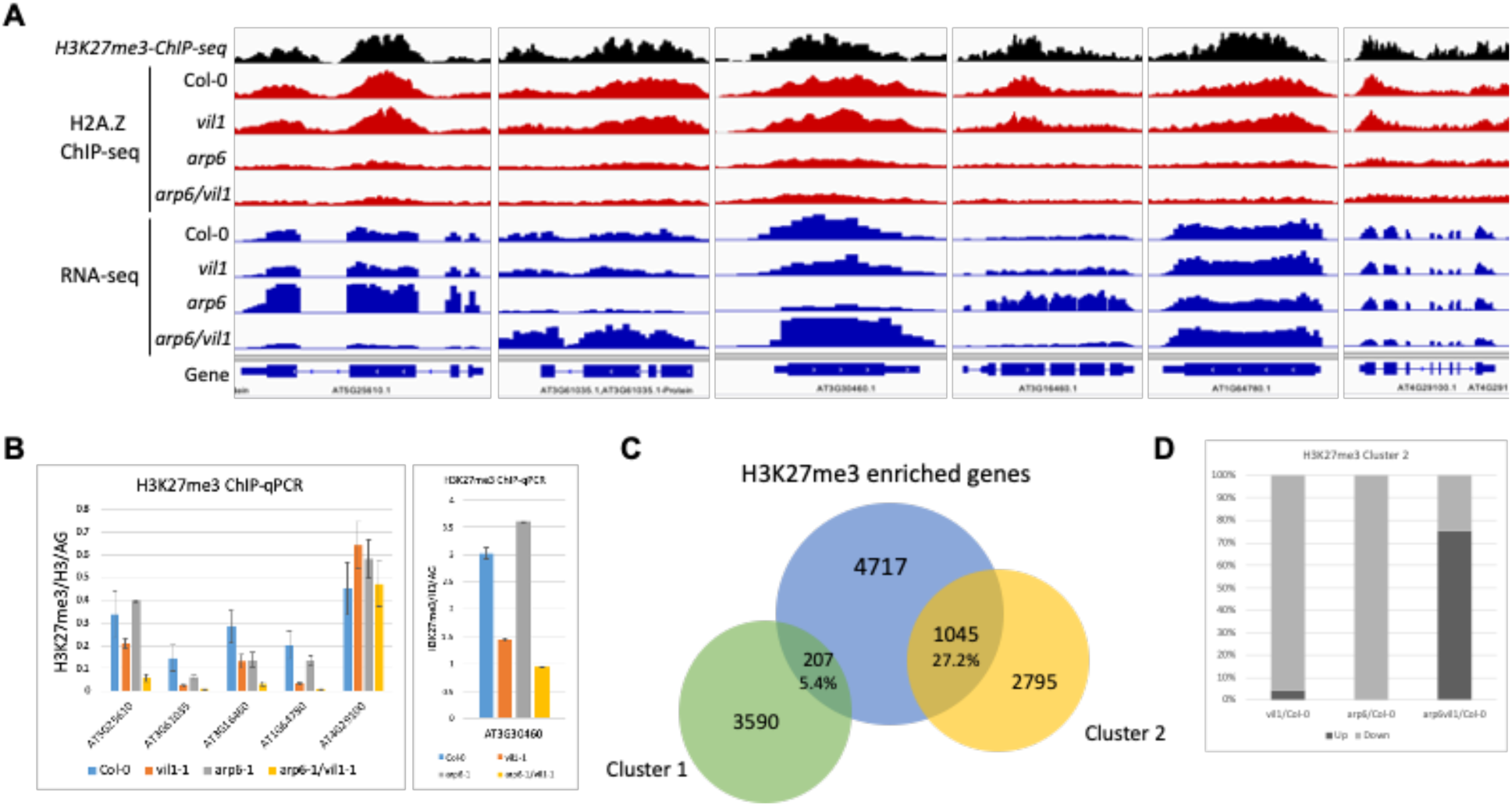
Antagonism between VIL1 and ARP6 in part can be explained by H3K27me3. **A**. Randomly selected transcriptionally reversed genes for ChIP-qPCR. **B**. H3K27me3 level normalized to H3 and Agamous gene. **C**. Overlap analysis of H3K27me3 enriched genes and cluster 1 and cluster 2 genes. **D**. Proportion of genes up and downregulated among 1045 genes which are common between H3K27me3 and cluster 2.

Our results clearly show that VIL1 plays an important role in warm temperature response in Arabidopsis through its function in imparting H3K27me3 and removal of H2A.Z. To support this, we also show a strong genetic interaction/antagonism between VIL1 and ARP6. This study provides a genetic basis of the interaction between chromatin modifications and histone variants to mediate responses to environmental changes.

## Discussion

In this study we reveal a new player in the high ambient temperature response. Role of PRC2 in vernalization in response to cold and in other plant developments has been widely studied and well established. Our findings provide new insight on the role of PRC2 associated protein during development at high temperatures. PRC2 plays an important role not only in the growth, development and differentiation ^34^ but also in response to stress ^35^. It is interesting to note that VIL1 was originally isolated as vernalization insensitive mutants which mediate cold temperature response. The same protein also plays role in the other spectrum of temperatures, high ambient temperature. Our result also implies that interaction between H2A.Z and H3K27me3/PRC2 dramatically affect the gene expression.

During this study, we found an antagonistic relationship between VIL1-mediated H3K27me3 and SWR1 complex-mediated H2AZ deposition. Broadly speaking, this reflects antagonism between chromatin modifications. A myriad of histone modifications coexist on the chromatin, studying how presence of one affect another is very important to understand how transcription is regulated in Eukaryotes. Extensive research in embryonic stem cells (ESCs) suggests that H2A.Z preferentially localizes to promoters of inactive genes occupied by the PRC2 ^36,37^. In addition, H2A.Z is also enriched at enhancers in ESCs and facilitates the binding of PRC2 and H3K4me3 and H3K27me3 modifications ^38^. Such enhancers have also been referred to as poised enhancers. These studies suggest that the interaction between H2A.Z and PRC2 might be much more complex and involve additional histone modifications. A previous study in Arabidopsis utilized pure bioinformatics approach to study the relationship between histone variant H2A.Z and histone modifications such as H3K27me3 and H3K4me3 ^39^. However, to this date there is no report showing genetic evidence of interaction between components of PRC2 and that of SWR1 complex in plants. To confirm that this antagonism is not an indirect effect of the large pleiotropic changes in *arp6* genetic background, we also generated *vil1/hta9/hta11* triple mutant. HTA8, HTA9 and HTA11 are three genes in Arabidopsis which code of histone variant H2A.Z. *hta9/hta11* double mutant phenocopy *arp6*. As expected, *vil1* suppresses the early flowering phenotype of hta9/hta11 and the overall phenotype of *vil1/hta9/hta11* triple mutant resembles more like *arp6vil1* (Supp Fig. 6). Wang et al. showed interaction among H2A.Z, H3.3 and PRC2-dependent H3K27me3 in mouse embryonic stem cells ^40^. They suggest that H3.3 counteracts the action of H2A.Z. It is possible that VIL1 or PRC2 indirectly affect the level of H3.3 in the Arabidopsis genome leading the antagonism between these modifications.

We ascribe the hyposensitivity of *vil1* at high ambient temperature to two main factors; 1) lack of H3K27me3, because *vil1* fails to downregulate the heat suppressed genes, and: 2) lack of H2A.Z eviction in *vil1* in from heat responsive genes. Given these two reasons, it is not surprising that *vil1* and *arp6* show a strong genetic interaction. It is likely that VIL1 or PRC2 by extension could be involved in eviction of H2A.Z in Arabidopsis. Although it is beyond the scope of this study, to address this possibility, the interaction between PRC2 associated proteins and INO80 complex needs to be addressed. INO80 is required for maintaining the boundary of H2A.Z at +1 nucleosome by evicting the ectopically deposited H2A.Z by SWR1 complex ^41^.

Our study enhances our understanding of how eukaryotes utilize epigenetic mechanisms under unfavorable environmental conditions. The insights developed from such research will lead us to develop better strategies to thrive in the era of climate change.

## Acknowledgements

The authors acknowledge the Texas Advanced Computing Center (TACC; http://www.tacc.utexas.edu) at The University of Texas at Austin for providing High

Performance Computing resources that have contributed to the research results reported within this paper. This work was supported by NIH R01GM100108 and NSF IOS 1656764 to S. S.

## Materials and methods

### Plant material and growth conditions

All experiments were done in Col-0 ecotype of Arabidopsis thaliana. The Arabidopsis T-DNA lines used in this research are: *vil1-1* (SALK_136506), *vil1-2* (SALK_140132), *arp6-1* (Garlic_599_G03), *hta9-1* (SALK_054814), *hta11-1* (SALK_017235). Plants were grown in long day (16-hour light, 8-hour dark) and short day (8-hour light, 16-hour dark) at 22°C or 27°C. VIL1::gVIL1-myc complementation line in *vil1-1* mutant background was used for VIL1 ChIP-qPCR. Experiment specific growth condition is also mentioned in the figure legend and/or results section. For heat treatment for RNA-seq, seedlings were germinated and grown on blue media (fertilizer) for 7 days at 22C followed by 9 days at 27°C. 22°C samples were continuously grown at 22°C for 14 days. For all the experiments samples were harvested at ZT-6.

### RNA-seq

Total RNA was extracted from seedlings (growth condition described above) by using TRIzol (Invitrogen) and treated with DNase I enzyme (Promega) to eliminate traces of genomic DNA. Sequencing libraries were prepared with 500 ng total RNA followed by NEBNext Poly(A) mRNA Magnetic Isolation and library preparation using NEB #E7420. Quality of library was assessed using Bioanalyzer (Agilent High Sensitivity DNA Assay). All the sequencing was performed on Illumina NextSeq 500 as 35×2 paired end.

For bioinformatics analysis reads generated by Illumina NextSeq 500 platform were checked for quality using FastQC. After quality assessment, reads were aligned on TAIR10 genome using HISAT2. SAM files generated from mapping were then converted into BAM files and sorted using Samtools. Bigwig files for IGV visualization were created using deepTools. For the gene counting bedtools program was used which generates raw count using BAM files. Raw count was then fed into R for differential gene expression analysis using edgeR and data visualization. For hierarchical clustering analysis only differentially expressed genes were used. Specific analysis detail is also provided in figure legend. Codes for the analysis can be provided upon request. Go term enrichment analysis was performed using Gene Ontology website http://geneontology.org/.

### H2A.Z ChIP-seq

Seedlings were crosslinked using 1% formaldehyde. ChIP procedure was followed as described in ^42^. Library was prepared from immunoprecipitated and input samples using NEBNext ChIP-Seq Library Prep Master Mix Set for illumine (E6200L). Quality of library was assessed using Bioanalyzer (Agilent High Sensitivity DNA Assay). For bioinformatics analysis reads generated by Illumina NextSeq 500 platform were checked for quality using FastQC. After quality assessment, reads were aligned on TAIR10 genome using Bowtie2. SAM files generated from mapping were then converted into BAM files and sorted using Samtools. Bigwig files for IGV visualization were created using deeptools. MACS2 broad peak call was used for peak calling (q <0.01). For differential binding analysis, DiffBind package was used in R and MA plots were generated using the same package. For H2A.Z enrichment analysis, deepTools package was used. First, computeMatrix was generated and then plotProfile function was used to generate the enrichment profiles.

### qRT-PCR

RNA was extracted using TRIzol reagent (Ambion). 1μg total RNA was used to synthesize cDNA. After DNaseI treatment to remove any DNA contamination, random primer mix (NEB #S1330S) and M-MLV Reverse transcriptase (Invitrogen #28025-013) were used for first strand synthesis. qRT-PCR was used to quantify the RNA prepared from transient expression experiments. AzuraQuant qPCR Master Mix (Azura Genomics) was used with initial incubation at 95°C for 2 min followed by 40 cycles of 95°C for 10 sec and 60°C for 30sec. Level of target RNA was calculated from the difference of threshold cycle (Ct) values between reference and target gene. PP2A gene was used as reference. List of primers used for qRT-PCR is provided in supplementary table 1.

### ChIP-qPCR

Seedlings were crosslinked using 1% formaldehyde. ChIP procedure was followed as described in ^42^. Antibodies used for ChIP were ab1791 (H3), ab5408 (Pol II S5), ab1220 (H3K9me2), ab47915 (H3 acetylation), ab8580 (H3K4me3), ab11946 (H3K4me2) and Millipore 07-449 (H3K27me3). For gVIL1-myc ChIP anti c-Myc antibody was used (Santa Cruz Biotechnology # 9E10). After ChIP, DNA was purified using Qiagen PCR purification kit (QIAquick 28106). AzuraQuant qPCR Master Mix (Azura Genomics) was used with initial incubation at 95°C for 2 min followed by 40 cycles of 95°C for 10 sec and 60°C for 30sec. For all histone ChIP-qPCR, H3 ChIP was used for normalization. List of primers used for ChIP-qPCR is provided in supplementary table 1.

## Supplementary information

Supplementary Figures S1 ∼ S7

Supplementary Tables S1∼S3

Primers used for qRT-PCR and ChIP-qPCR (supplementary table 1)

H2A.Z ChIP-seq peak information (MACS2) (Supplementary table 2)

Raw read count and RPKM normalized RNA-seq count (Supplementary table 3)

