## Supplementary Figures for "VIL1, a Polycomb-associated protein, modulates high ambient temperature response via H3K27me3 and H2A.Z in *Arabidopsis thaliana*"

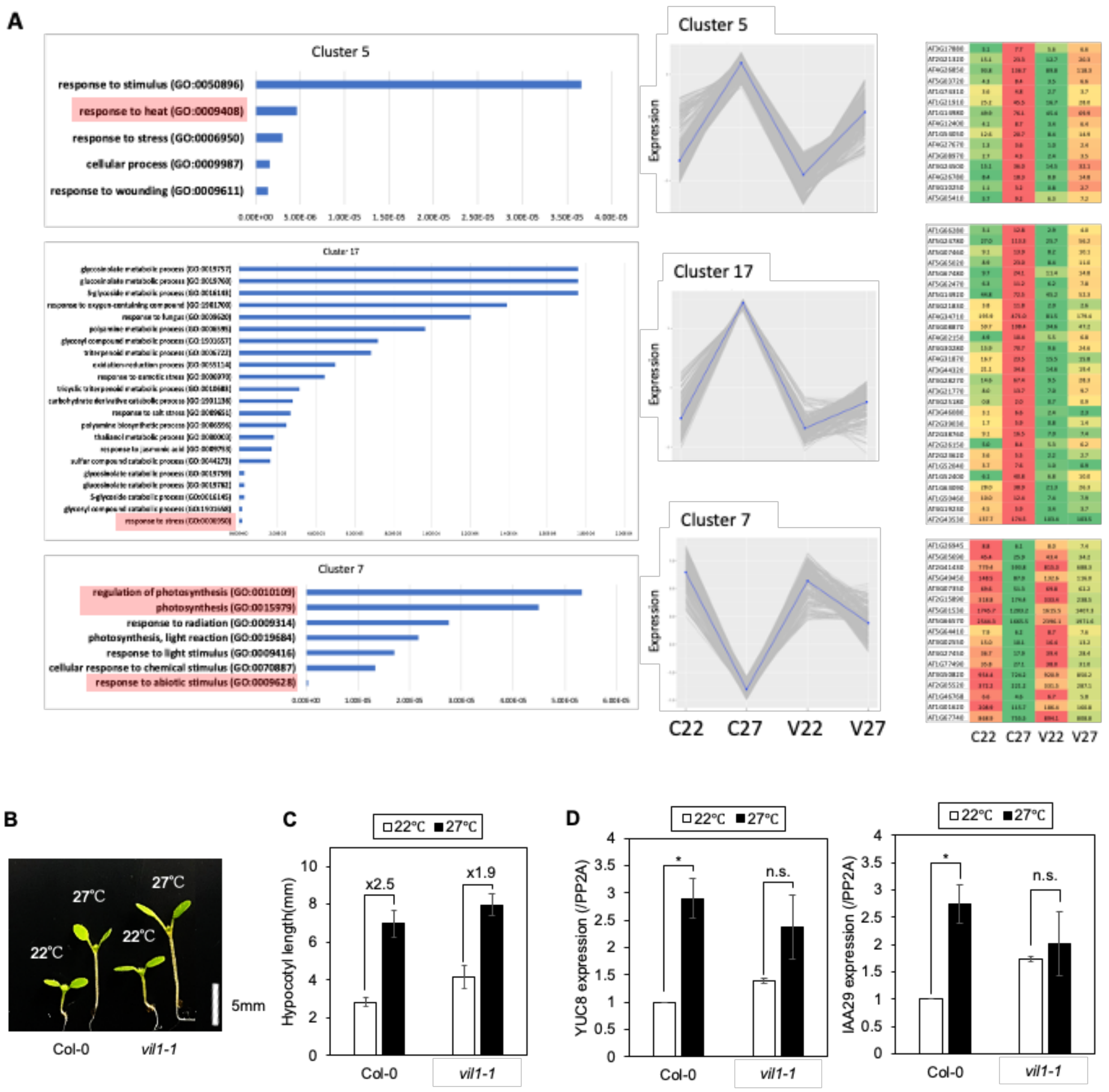


**Supplementary figure 1. GO term analysis and hypocotyl length phenotype**

**A.** GO term analysis on clusters 5, 17 and 7 on left. The panel in the middle shows expression pattern in each cluster. While induction of genes in clusters 5 and 17 is poor in case of vil1 compared to Col-0 at 27°C, cluster 7 genes are not downregulated in *vil1* as much as WT in response to high temperature. Panel in the right shows color coded average RPKM values of randomly selected genes from each cluster. **B.** Hypocotyl length phenotype of Col-0 and *vil1-1* at 22°C and 27°C. Seedlings were grown at 22°C for four days and then transferred to 27°C for three days to check hypocotyl length and RNA analysis. **C.** Quantification of hypocotyl length showing fold change in *vil1-1* when moved from 22°C to 27°C. **D.** RNA analysis of Auxin signaling downstream genes which are required for hypocotyl length elongation. While YUC8 shows only 2-fold change in expression from 22°C to 27°C in *vil1*, in Col-0, the fold change is 3.2-fold. IAA29 is not significantly induced in *vil1-1*. * ; $p$<0.05, n.s.; not significant

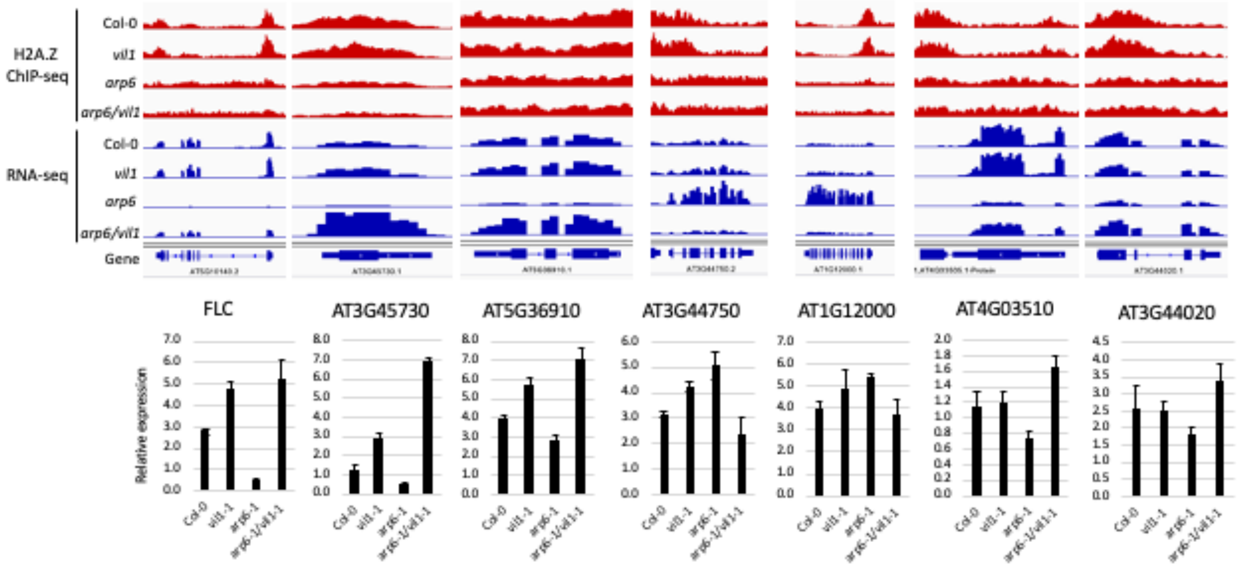


**Supplementary figure 2. qRT-PCR confirmation of RNA-seq result**

Randomly selected transcriptionally reversed genes (shown as IGV snapshots) were used to confirm the RNA-seq result. Expression of genes relative to PP2A shown below.

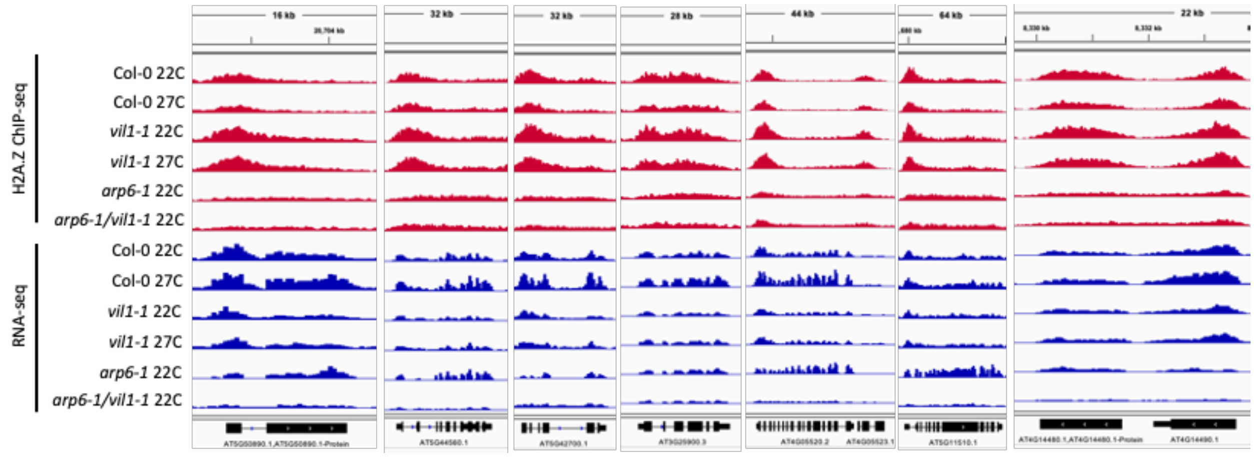


**Supplementary figure 3.** IGV snapshots with H2A.Z ChIP-seq and RNA-seq data of randomly selected genes which fail to evict H2A.Z in vil1 in response to high temperature.

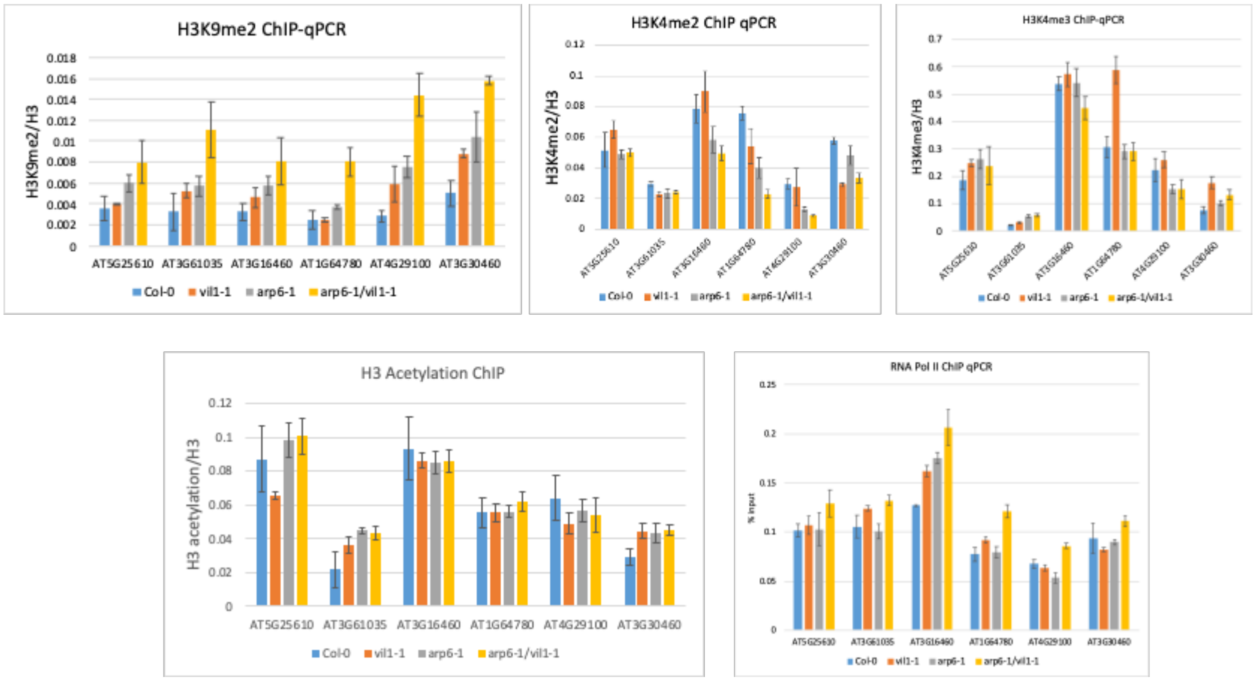


**Supplementary figure 4. A number of histone modifications remain unchanged in mutants**

ChIP-qPCR to check relative enrichment of histone modifications H3K9me2, H3K4me2, H3K4me3, H3 acetylation and RNA pol II (S5).

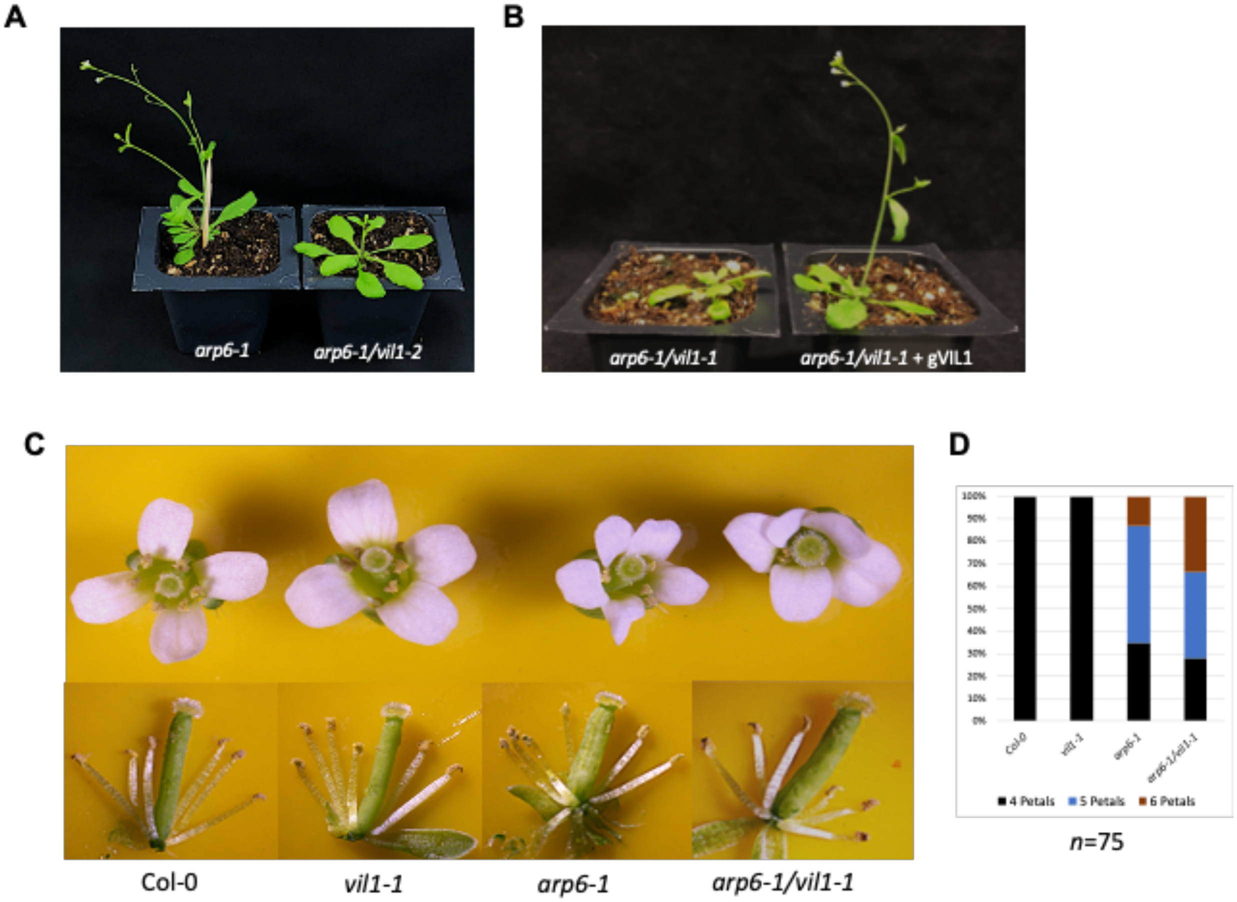


**Supplementary figure 5. A.** Genetic interaction between arp6 and *vil1* was validated with *vil1-2* allele. **B.** *arp6-1* single mutant phenotype reappeared when *arp6-1/vil1-1* double mutant was crossed with a VIL1 complementation line with genomic VIL1 driven by native promoter. **C.** Flower phenotype of Col-0, *vil1-1*, *arp6-1* and *arp6-1/vil1-1*. Major difference in the flower is number of petals in *arp6* and *arp6vil1*. **D.** Quantification of petal number.

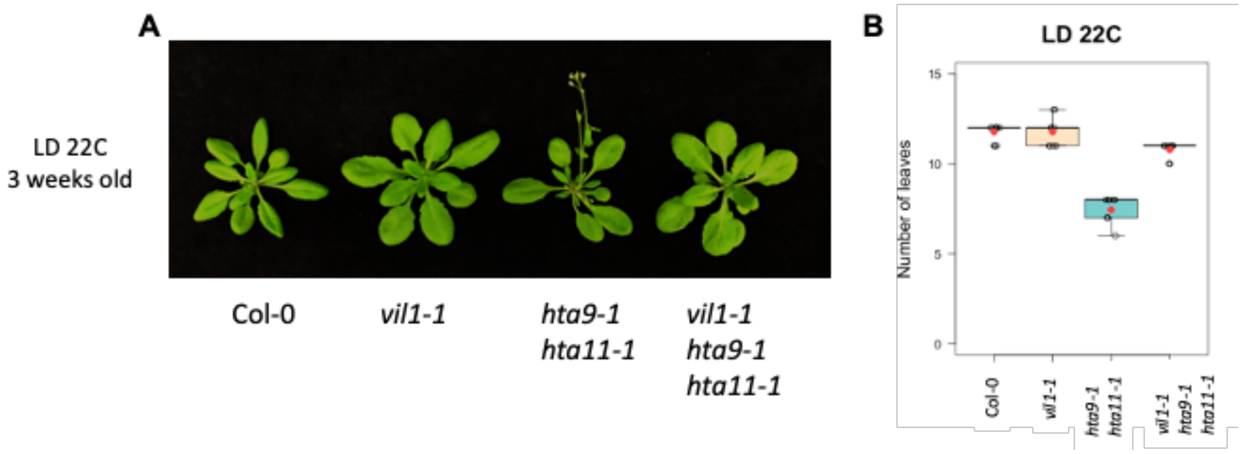


**Supplementary figure 6. A.** *vil1-1/hta9-1/hta11-1* shows same phenotype as *vil1-1/arp6-1.* Plants grown in long day at 22°C. **B.** Flowering time data shown in terms of number of leaves at the time of flowering. Red dot in the box plot indicate mean number of leaves.

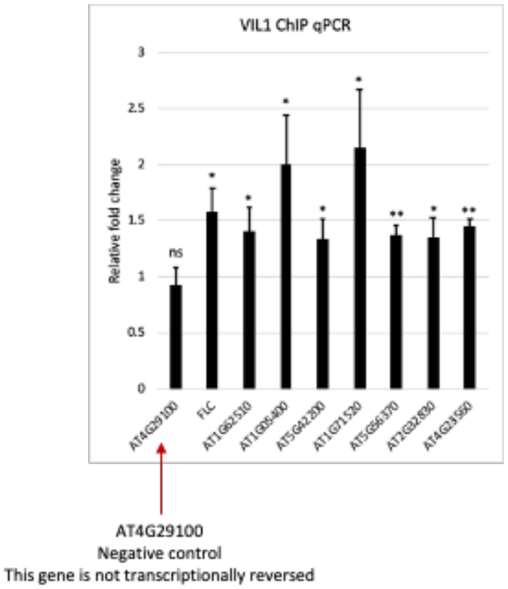


**Supplementary figure 7.** VIL1 ChIP-qPCR shows enrichment of VIL1 on transcriptionally reversed loci. Genes for ChIP-qPCR were chosen randomly from among 1045 genes which show overlap between cluster 2 and H3K27me3 as shown in figure 7C. AT4G29100 is not transcriptionally reversed in *arp6/vil1* and shows no VIL1 enrichment. Relative fold change was calculated compared to 5s rDNA. Statistical significance was determined using T-Test. (ns - Not Significant)

* T-Test P value <0.05

** T-Test P value <0.01
